# Integrating Equation Coding with Residual Networks for Efficient ODE Approximation in Biological Research

**DOI:** 10.1101/2024.01.01.573840

**Authors:** Ziyue Yi

## Abstract

Traditional biological research heavily relies on experimental methodologies, which often lead to inefficiencies in communication and knowledge transfer. These methodologies frequently result in redundant trial-and-error experiments, posing challenges in standardizing results and impeding rapid knowledge dissemination. The complexity of biological systems and the sheer volume of experimental data generate a demand for precise mathematical models like Ordinary Differential Equations (ODEs) to articulate biological interactions. However, practical implementation of ODE-based models is hampered by their need for data curation, making them less feasible for everyday research applications. To address these limitations, we introduce LazyNet, a novel computational model that employs logarithmic and exponential functions embedded within a Residual Network (ResNet) architecture to approximate ODEs. LazyNet simplifies complex mathematical operations, reducing the dependency on large datasets and extensive computational resources. This study tests LazyNet across various biological scenarios, including HIV dynamics, gene regulatory networks, and mass spectrometry analysis of small molecules. Our results demonstrate that LazyNet can efficiently assimilate and predict complex biological processes, significantly enhancing model training speed and reducing the reliance on empirical experimentation. Therefore, a promising tool for advancing biological research and fostering more accurate and efficient scientific explorations.

## Introduction

Traditionally, biology has been dominated by experimental approaches, relying on descriptive methodologies for both communication and academic research. This heavy reliance has led to challenges in standardizing communication and transferring knowledge, often resulting in costly and redundant trial-and-error experiments. To mitigate these challenges, there has been a significant push to integrate mathematical modeling into biological research, particularly through the use of Ordinary Differential Equations (ODEs) to articulate complex biological phenomena.

ODEs conceptualize variables with their derivatives structured as equations that incorporate biological interactions. This modeling facilitates detailed predictions of biosystem states, allowing common experimental interventions such as inhibition or overexpression to be modeled as modifications to initial conditions.

Despite their utility, ODEs encounter substantial challenges in modeling biological systems both accurately and efficiently. A pivotal issue is the integration of experimental data into ODE models, a process that traditionally requires extensive input from biomathematicians in areas such as text mining, data curation, and empirical equation development.^1^ These models often suffer from a lack of standardization and are slow to integrate new discoveries, thereby constraining their utility in biological research.

Recent advancements in differential equation modeling within physics and mathematics have significantly influenced the field of biological modeling. The sheer diversity and volume of biological data make it an excellent candidate for these advanced methodologies. Beginning in 2009, pioneers such as Michael Schmidt and Hod Lipson utilized computational techniques to derive equations directly from experimental data.^2^ This trend was advanced in 2018 when Ricky Chen incorporated maximum likelihood estimation with ODE solvers into neural networks, further refined by John Qin’s application of these techniques within the Residual Network (ResNet) framework and Samuel Kim’s introduction of symbolic dictionaries for equation discovery.^3–5^ Chen, Liu, and Sun extended this line of work by integrating autograd differentials into the networks, enhancing the physics-informed learning from sparse data.^6^ Although categorized under symbolic regression, these methods demand substantial datasets even for relatively simple differential equation systems and significant computational resources, which restrict their practical application in biological research.

In an alternative approach, Alex Liu introduced the Kolmogorov–Arnold Networks (KAN) that dynamically encode equations using spline functions.^7^ While this method reduces layer size and parameter count, it remains computationally demanding and time intensive.

To address these biological and computational challenges, we developed LazyNet. This innovative model uses logarithmic and exponential functions embedded within the weights of a ResNet architecture to approximate ODEs through Euler method. Supporting a broad spectrum of mathematical operations — including addition, subtraction, multiplication, division, exponentiation, and Taylor series expansions— LazyNet simplifies the complexities associated with spline functions and sidesteps the limitations of symbolic regression methods. Requiring fewer resources, LazyNet operates effectively with datasets under 30,000 samples on standard computation clusters.

In this study, LazyNet has been rigorously tested across various biological applications, including the dynamics of HIV populations, gene regulatory networks (GRN), and the analysis of small molecule mass spectrometry (MS).^8–11^ These tests have confirmed that models trained with LazyNet can effectively assimilate and predict biological processes, demonstrating its capability and practicality in constructing ODE models for biosystems.

## Results

Consider a biosystem where the state of the system at any given point is dependent on its preceding state, forming a continuous sequence defined by a differential equation that reflects the intrinsic nature of the system’s interactions.

Let the sequence of states be represented as ***f1, f2, f3, …***, where ***f1*** denotes the initial state of the system, ***f2*** the state at the next time step, and so forth. Each state in this sequence evolves according to the function ***F***, which is consistent across the system and encapsulates the biological interactions. Using Euler’s method for the approximation of these dynamics, the progression of states can be mathematically modeled by the equations:

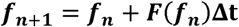

where Δ***t*** denotes the time step increment, and ***f_n+1_*** represents the state at the subsequent time point. Each state ***f_n_***encompasses the conditions of variables such as ***x,y,z,***…, allowing for an iterative computation where each subsequent state is derived directly from its predecessor.

System ***F***, plays a critical role in the sequential transition from state ***f_n_*** to ***f_n+1_*** within biological models. It encapsulates the inherent biological interactions and mechanisms, serving as a fundamental component in understanding the system’s dynamics, such as gene X influences over gene Y. The concept of data-driven discovery of ODE governing equations aims to automate the formulation of ***F*** using large datasets. In LazyNet, this formulation is achieved by:

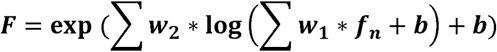

From the entire network layout, the logarithm and exponential layers are served as residue block. (Figure 1) At the end of network, an addition layer integrates these transformations with the current state and output the next state, meanwhile complete Residual Networks (ResNet) and Euler structure.

**Figure 1.**
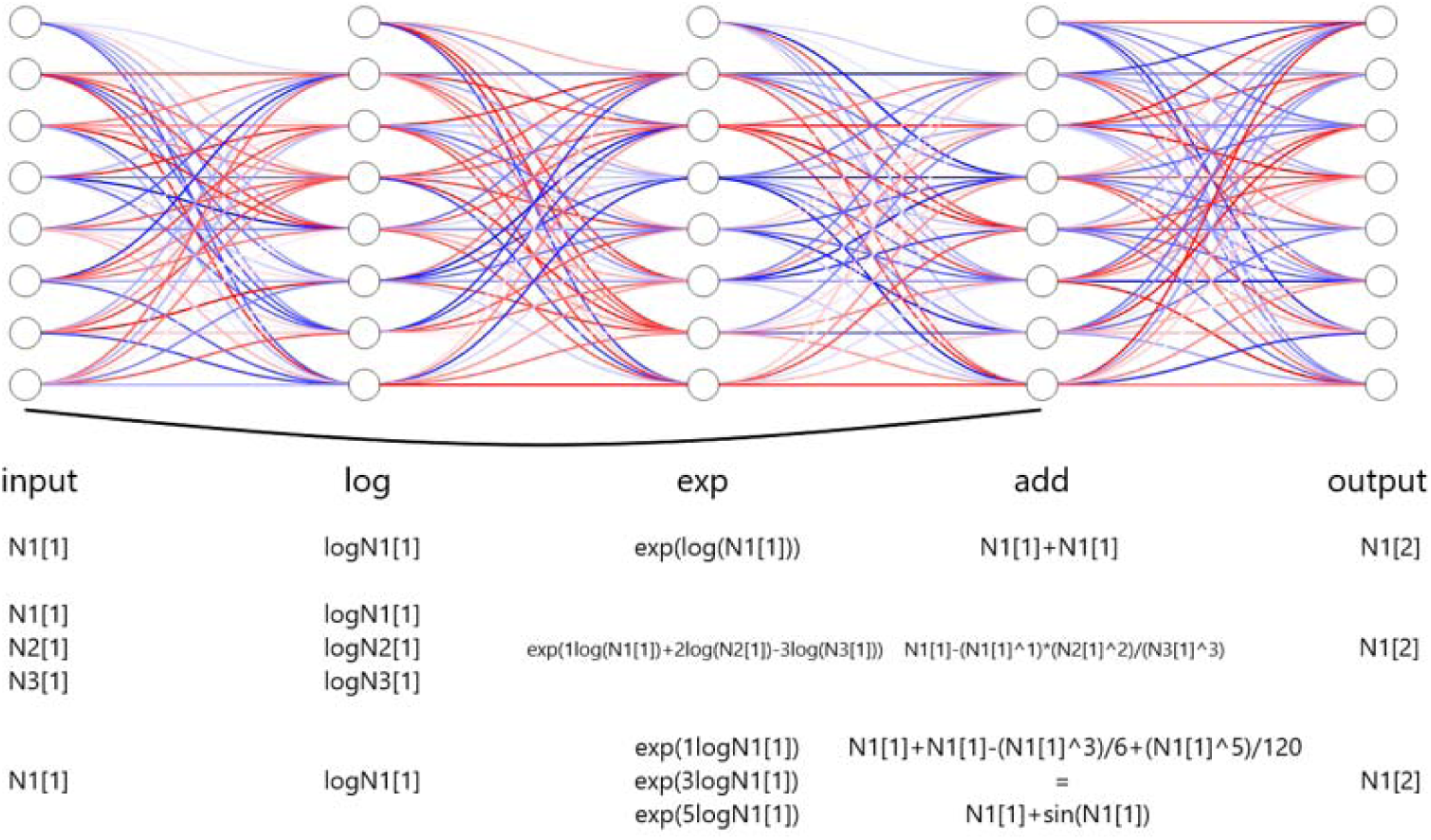
Schematic architecture of the framework of LazyNet. LazyNet unique design to approximate ODEs within a ResNet framework using logarithmic and exponential transformations. The diagram highlights the main components of LazyNet, including its dynamic weight adjustment capabilities, simplified mathematical operations through log and exp functions, and the integration of these elements into the output.

This architecture is characterized by its dynamic weight adjustment feature, where both weights (***w***) and biases (***b***) are learnable parameters, finely tuned during the training process. LazyNet utilizes these adjustable parameters to conduct key operations on inputs, such as addition, multiplication, and Taylor series expansions. In some other words, LazyNet employs logarithmic and exponential functions to encode operations. This approach allows the weights to serve multiple functional roles, significantly streamlining the network complexity.

Contrary to traditional methods that depend on extensive symbolic operation libraries or numerous parameters for defining linear operations like spline functions, LazyNet simplifies these processes. Additionally, by integrating L1 regularization, LazyNet is designed to provide the most parsimonious approximation of biological systems. For example, if two polynomial terms are sufficient to approximate a sine function, LazyNet will avoid using an unnecessary third term, showcasing its “lazy” yet effective design principle.

LazyNet utilizes the Adam optimizer and employs Huber Loss as its loss function. By consider the batch issues, the total loss function:

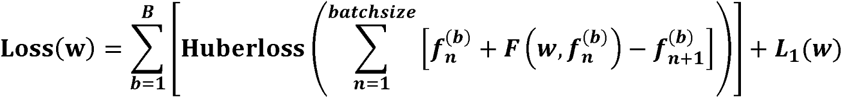

Where **w** is the learnable weight, **b** is the batches to the last batch **B**, the training is base on each point **n** within batch **b**, and **L_1_** is the L1 regularization.

### LazyNet’s Estimation of HIV Strain ODE System

To assess the efficacy of LazyNet, we utilized a simplified biological model of HIV population dynamics articulated through 19 ordinary differential equations (ODEs).^8^ These equations, selected for their simplicity and absence of trigonometric or complex functions, aim to reduce baseline complexity.

Considering the exploratory nature of this research and the unpredictable iterations required for validating LazyNet, we synthesized a dataset using MATLAB that mirrors typical variances found in virological studies, influenced by different initial infection populations. We began with 100 samples, each subjected to a unique random alteration in initial conditions. As the scenarios grew in complexity, the sample size was meticulously reduced to 894, analyzed across 500 temporal points (Figure 2A). For the testing phase, we crafted a dataset with random modifications to all initial values, through which we assessed LazyNet’s performance across 50 distinct test datasets (Figure 2B).

**Figure 2.**
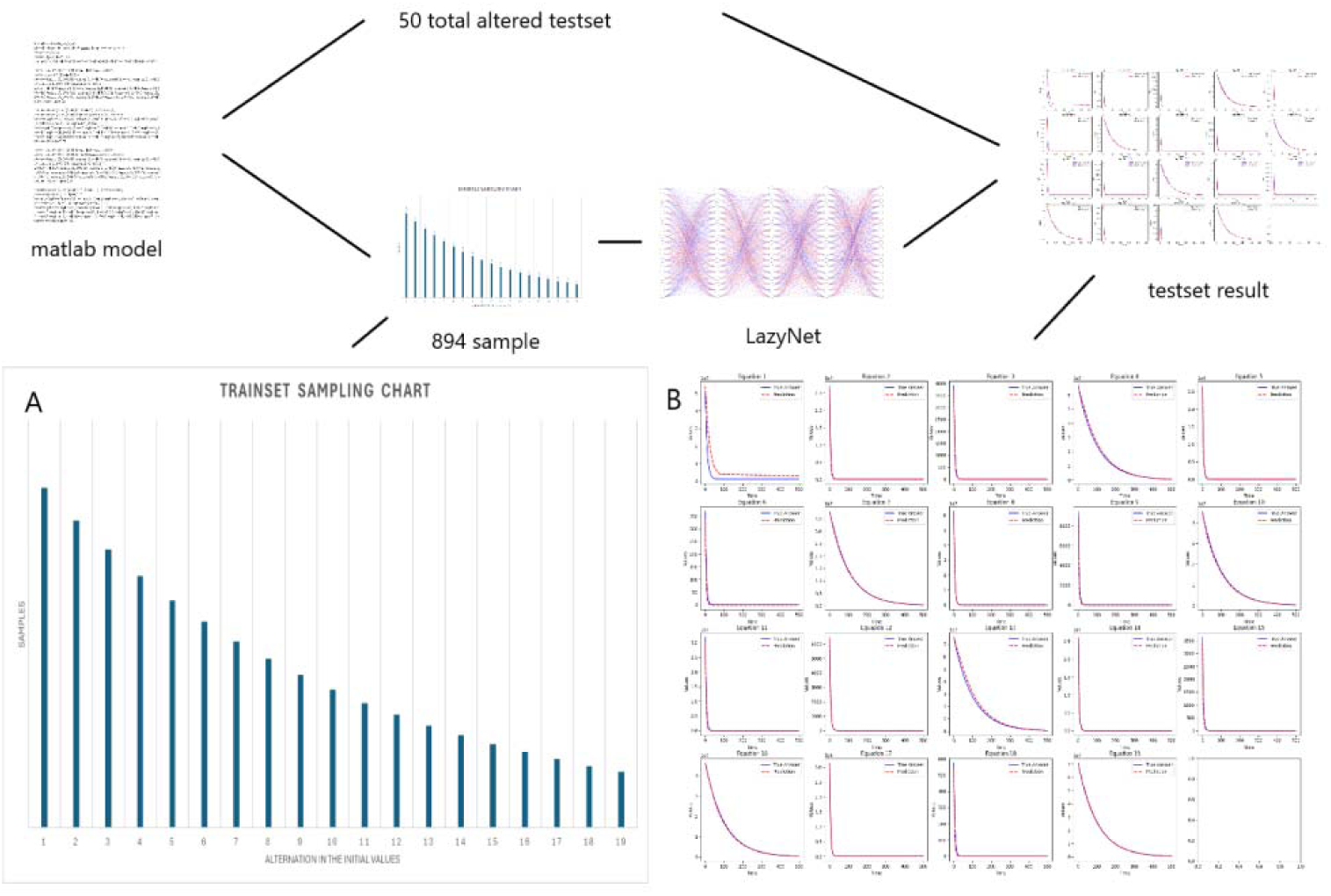
Workflow and Results from LazyNet’s Evaluation on HIV Dynamics. (A) Trainset Sampling Chart showing the distribution of 894 samples each uniquely altered in initial conditions. (B) Performance on 50 altered test datasets displayed through plots that compare LazyNet’s predictions (red lines) against expected outcomes based on the MATLAB model (blue lines). The result plot represents one of the test case, and each subplot represents one strain.

The initial results are encouraging (Supplement S2). LazyNet demonstrated a robust capacity to emulate the dynamic behaviors delineated by the ODE model, thereby validating its accuracy and fidelity. Upon preparing the synthetic dataset, we fine-tuned LazyNet to identify optimal training parameters within a few days—a notable enhancement over conventional manual modeling approaches. This improvement in training efficiency significantly streamlines the modeling process.

However, a limitation was observed in the dynamic range of the system. The generated dynamics were excessively stable, exhibiting very similar patterns despite variations in numerical values. This homogeneity points to a potential deficiency in LazyNet’s ability to accommodate more complex and dynamically varied scenarios. This finding underscores the need for further investigations with more intricate ODE systems to thoroughly evaluate LazyNet’s capabilities in modeling complex dynamic behaviors.

### Adapting to Periodic Relationships in Gene Regulatory Networks (GRN)

To explore LazyNet’s aptitude in managing intricate biological systems and interpreting periodic relationships, we applied it to a Gene Regulatory Network (GRN) based on a model described in established literature.^9^ Omics-related data, which are pivotal in various biological studies, present a vast application scope for LazyNet. This is especially pertinent in experiments like gene knockouts or overexpression aimed at predicting the outcomes of physical experiments.

Our investigation employed a GRN model constituted by seven ordinary differential equations (ODEs) that mimic periodic biological functions similar to sine and cosine waves.

Synthetic data were generated using MATLAB, adhering to the methodological protocols previously described. The study commenced with 100 samples, each subjected to a random modification in initial conditions. As the complexity of the experiments increased, the sample size was judiciously reduced, culminating in 529 data points per set across 500 time points over 50 sets, totaling 26,450 samples (Figure 3A). The test set comprised 50 samples with altered initial conditions, presenting the model with unforeseen scenarios (Figure 3B, Supplement S3). This design mirrors typical gene manipulation scenarios, such as knockouts and overexpression, providing a robust test of LazyNet’s capabilities in a controlled setting.

**Figure 3.**
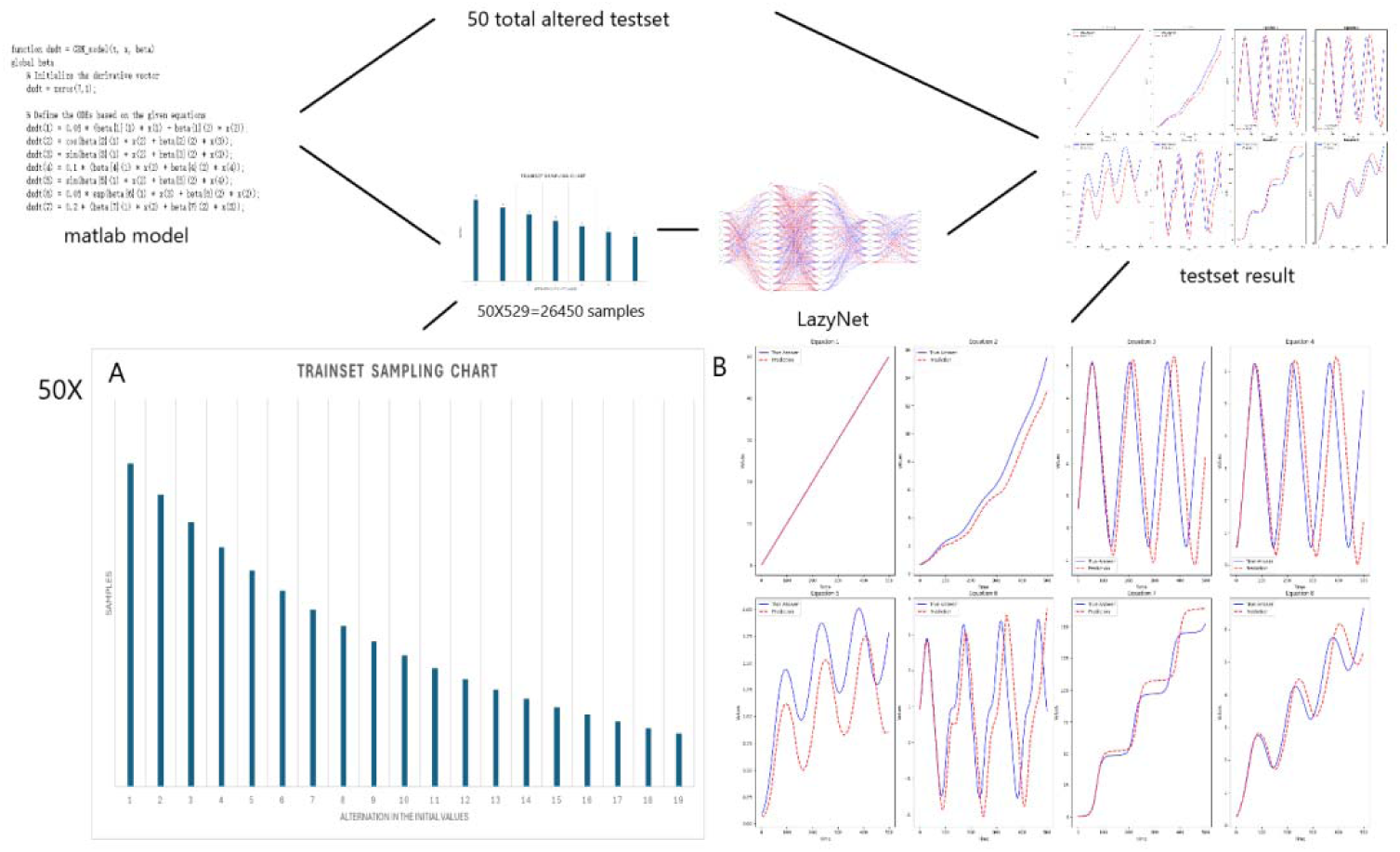
Workflow and Results from LazyNet’s Evaluation on Gene Regulatory Networks (GRN) (A) Trainset Sampling Chart illustrating the distribution of 26,450 samples across 500 time points over 50 sets, each starting from 100 sample s uniquely altered in initial conditions. (B) Performance on test datasets displayed through plots that compare LazyNet’s predictions (red lines) against expected outcomes based on the MATLAB model (blue lines). The result plot represents one of the test case, and each subplot represents one strain.

The test involved 50 randomly selected samples, all of which were predicted with remarkable accuracy (Supplement S3). These results not only confirm LazyNet’s precision but also underscore its vast potential in omics research. It enables researchers to predict the outcomes of physical experiments and pre-experimental planning. This capability is particularly beneficial in the fields of life sciences research and synthetic biology, where cost efficiency and experimental precision are paramount.

### Real-World Data in Mass Spectrometry and Synthetic Biology

Building on its previous successes, this study aimed to explore the significant role of LazyNet in the real-world application of mass spectrometry data analysis. LazyNet was employed to evaluate mass spectrometry data from eight genetically modified E. coli strains, which were engineered to synthesize distinct biochemical products. Despite facing substantial challenges, including the limited availability of datasets (only nine out of an expected 100) and the lack of detailed formatting in the available proteomics data, this scenario provided an ideal opportunity to test LazyNet in extreme conditions, thereby enhancing our understanding of its capabilities in practical applications.

To adapt to these constraints, the study was restructured to focus on using LazyNet to predict phenotypic behaviors across strains, employing only the available metabolomics data. The datasets contained data on 79 small molecules, and the network was designed to accommodate 80 inputs, including one additional variable to account for time. To refine our analysis and improve the resolution of the datasets, each was expanded to 500 data points using the Savitzky-Golay filter.

The results of our study, while promising, also identified areas for improvement. Of the 79 small molecules analyzed, 27 aligned with expected dynamic trends. (Figure 4A, Supplement S4). Furthermore, 23 of these molecules achieved a standard scaled RMSE lower than 1, demonstrating high predictive accuracy (Figure 4B). Consider some dynamically accurate results are not high in RMSE, the useful predictions are more than 30. The overall predictive accuracy was not entirely satisfactory, indicating that model performance may be constrained by the limited training data available. On the other hand, this scenario is particularly relevant in practical settings where researchers often have to make the most of low-quality or limited datasets. LazyNet can extract actionable insights from such data, enabling data recycle for valuable predictions.

**Figure 4.**
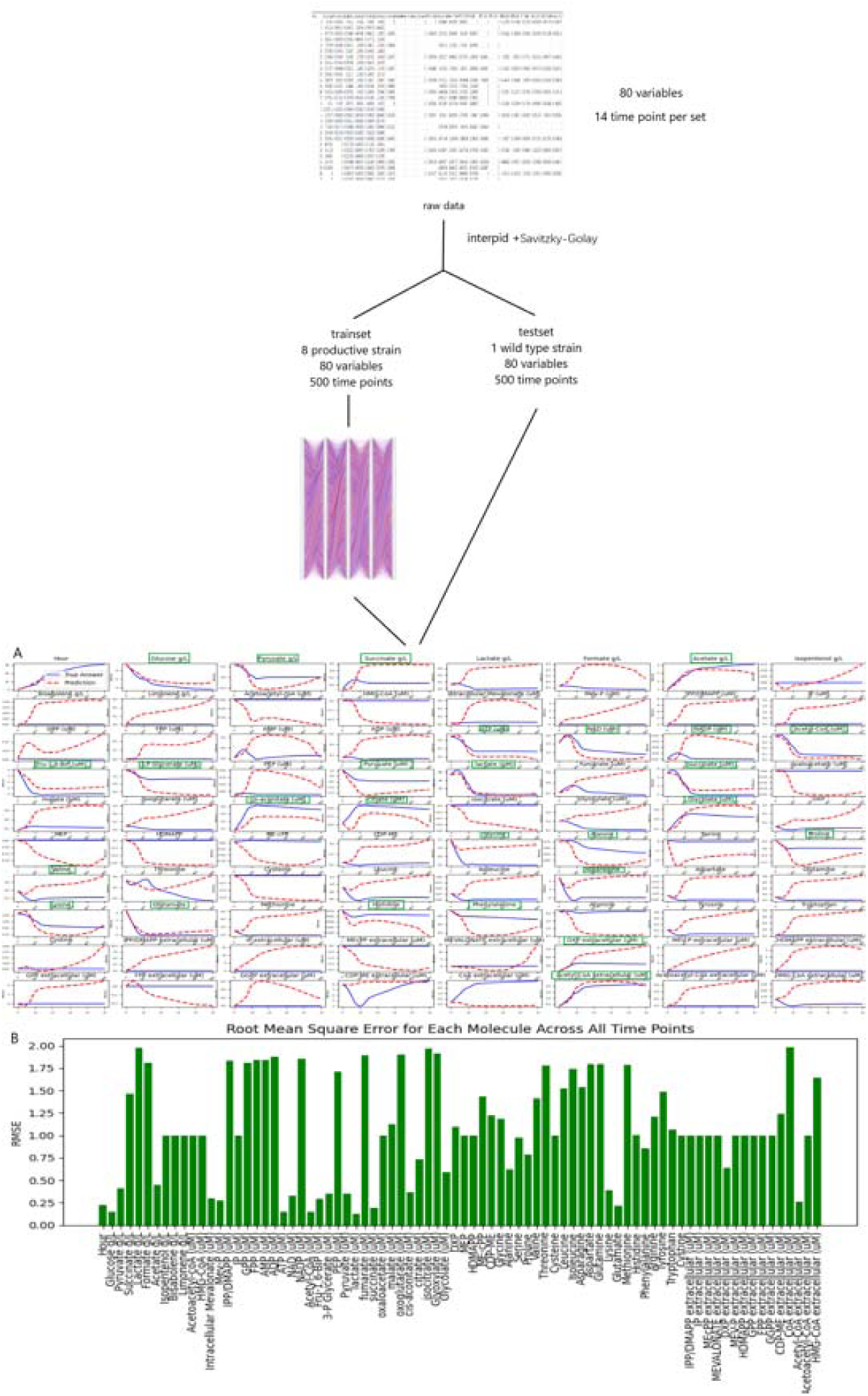
Data Processing and Predictive Analysis of E. coli Strains Mass Spectrometry. The workflow begins with raw data, initially composed of 80 variables across 14 time points per set, processed using the interp1d function and Savitzky-Golay filtering to enhance data resolution. This refined data comprises the trainset and testset; the trainset covers eight productive strains, and the testset covers one wild type strain. (A) depict LazyNet’s predictive performance across various biochemical molecules, comparing LazyNet’s forecasts (red lines) to actual data (blue lines), and dynamically correct predictions (green rectangle). (B) displays the Root Mean Square Error (RMSE) for each molecule across all time points.

A comparative analysis with the original study, which employed TPOT and reported a min-max scaled RMSE of 0.02 (standard deviation ∼0.018) across ten training datasets, helps contextualize LazyNet’s performance. Our study achieved a scaled RMSE of 0.042, which is commendable given that our analysis was conducted with fewer datasets and was limited to metabolomics data, unlike the original study that benefited from a richer set of both proteomics and metabolomics data.

In this study, LazyNet achieved acceptable results under stringent conditions characterized by limited datasets and non-optimal data quality. The performance of LazyNet is commendable when compared to that of TPOT, especially considering it utilized significantly fewer training datasets. Notably, LazyNet was not only able to extract meaningful insights from the data but also provided predictions that could be utilized for future research endeavors. These findings underscore the applicability of LazyNet in real-world scenarios, particularly in the fields of mass spectrometry data analysis and biosynthesis outcome prediction. The ability of LazyNet to yield practical results from limited data highlights its potential as a valuable tool in accelerating scientific discovery and enhancing decision-making processes in experimental biology.

### Integrating LazyNet with scRNA-seq Data for Enhanced Trajectory Analysis

Single-cell RNA sequencing (scRNA-seq) represents one of the most advanced experimental techniques in biological research. A unique analysis enabled by scRNA-seq is RNA velocity, which uses RNA splicing dynamics to assign pseudo-time to cell states. This pseudo-time provides temporal information that is often scarce in other areas of biological research. The incorporation of LazyNet with RNA velocity analysis offers a novel approach to scRNA-seq data, enabling an unprecedented style of analysis. This integration provokes significant interest in employing LazyNet to harness the full potential of temporal data provided by RNA velocity, facilitating deeper insights into cellular dynamics.

In previous study, RNA velocity provided by ScVelo indicating possible cyclic behaviors in monocytes—highlighted within a red rectangle in Figure 5A.^10,12^ This behavior has been a topic of debate in earlier research focusing on the dominance of Ticam2 and Myd88 pathways. The sequencing data illustrated in Figure 5B captured a Ticam2-dominated phase (black arrow), where monocytes cycle back to a resting state following PBS exposure. However, activation of monocytes and related pathways, potentially involving Myd88, was not evident (red arrow).

**Figure 5.**
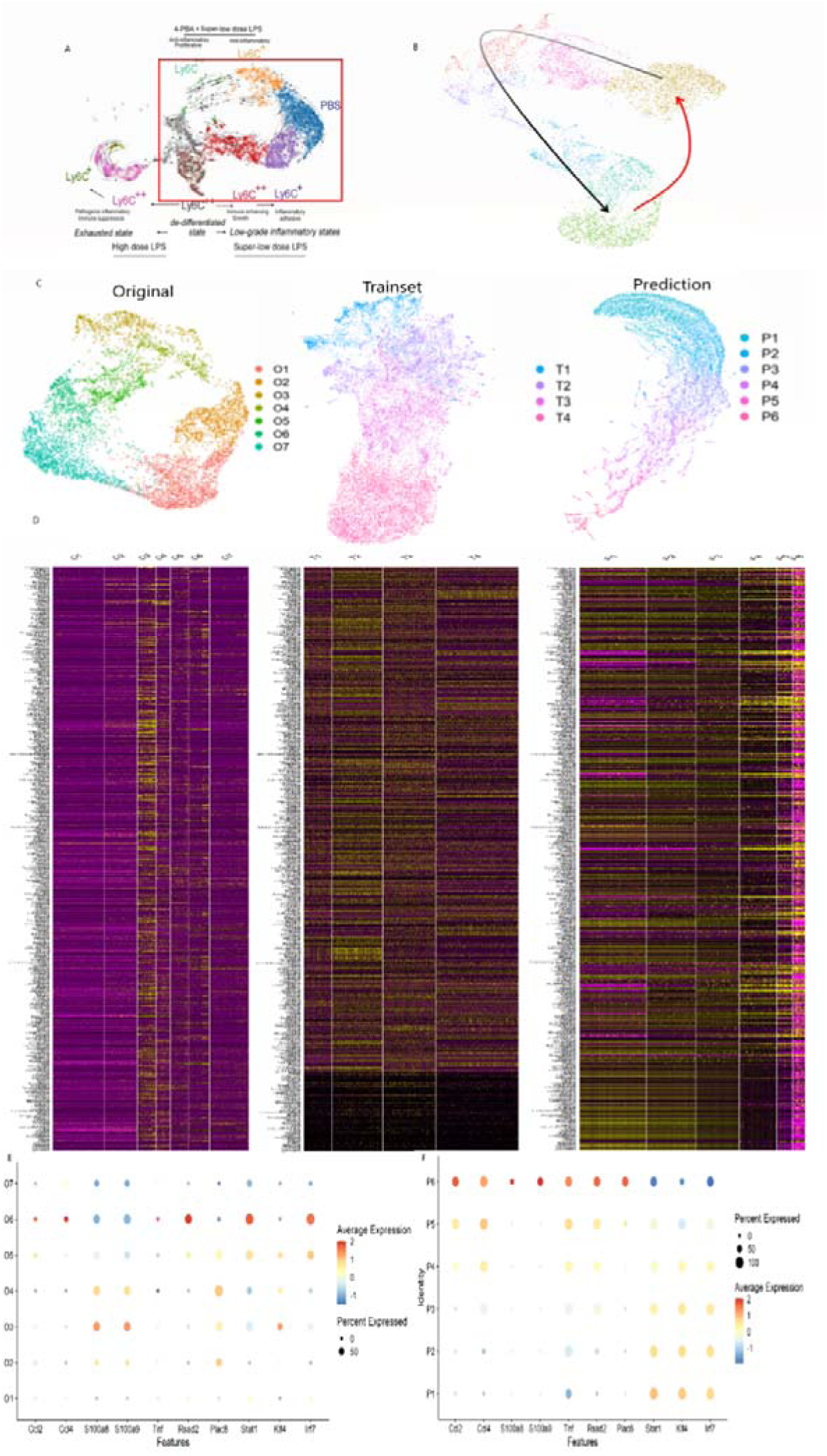
Comprehensive Analysis of Monocyte Dynamics Using LazyNet and ScVelo. (A) UMAP visualization highlighting potential cyclic behaviors in monocytes, with the red rectangle indicating areas of interest. (B) Ticam2 pathway (black arrows) denote ScVelo and LazyNet trajectories (red arrow), respectively. (C) Comparison of UMAP clusters between Original data, Trainset, and LazyNet Predictions, delineating shifts in cell state dynamics. (D) Heatmaps of gene expression across original, trainset, and prediction datasets, revealing gene regulatory patterns and pathway activations. (E) Dot plot representing differential expression levels in original data. (F) Dot plot representing differential expression levels in prediction.

LazyNet was utilized to extend the RNA velocity trajectory, facilitating an in-depth examination of the directionality and continuity of RNA velocity in monocyte populations. This approach enabled us to expand upon the foundational data, exploring cellular dynamics with greater precision.

Seurat employed to analyze 717 genes identified by principal component analysis (PCA), which facilitated the creation of a robust Uniform Manifold Approximation and Projection (UMAP).^13^ The dataset was expanded to include 140 data points using the Savitzky-Golay filter for smoothing. The UMAPs for the ***original data***, ***trainset***, and ***predictions*** are depicted (Figure 5C and Supplement S5), with the cluster names indicating the timeline. Contrary to initial expectations, the anticipated cyclic behavior in monocytes was not observed in the predictions. Extending this analysis further is deemed inappropriate as it would merely propagate the current dynamics without adding substantive insights, and could likely introduce errors.

In the heatmap (Figure 5D), the overall trend shows a decrease in gene expression over time, with clusters P5 and P6 exhibiting distinct patterns compared to the others. The trainset reflects the original data’s trends with a lesser decrease across all genes, while the prediction amplifies and extends this tendency, particularly in the lower genes. The dotplot (Figure 5E) reveals a shift from moderate to completely negative trends in the original data, a pattern that is mirrored from P1 to P4 in the predictions (Figure 5F). This observation supports the trends and cluster assignments.

Once the clusters are confirmed, further analysis of clusters P5 and P6 can be undertaken. Previous studies that focused on the dominance of the Ticam2 and Myd88 pathways described a trajectory from Ticam2 activation to PBS exposure. It is hypothesized that the other side of the PBS (Figure 5B, red arrow) is dominated by the Myd88 pathway.

Clusters P5 and P6 in the prediction were analyzed for differential gene expression compared to other clusters (Supplement S5). The dot plot indicates a decrease in the Ticam2 pathway and strong Myd88 signals (Figure 5F). In the prediction, genes like Ccl2, Ccl4, S100a8, S100a9, Tnf, Rsad2, and Plac8 are strong indicators of Myd88 pathway activity, with their upregulation indicating elevated activity.^14–20^ Conversely, genes typically associated with the Ticam2 pathway, such as Stat1, Irf7, and Klf4, were downregulated.^21–22^ To complete the monocyte dynamics theory, an experiment focused on the Myd88 pathway immediately post-LPS exposure should be prepared. The existing literature and physical experiments also support MyD88 activation immediately following LPS exposure, with the MyD88 pathway dominating.^23–24^ By knowing the potential cell state and reference literature, the experiment plan can save costs.

Although the anticipated cyclic behavior in monocytes was not observed, the study still yielded profound insights. The immediate dominance of the MyD88 pathway following LPS exposure was clearly supported by the data, suggesting a potential transition to a Ticam2-dominant state thereafter. This indicates a hypothetical dynamic interaction where cells may toggle between the Ticam2 and MyD88 pathways under specific conditions. These insights, easily derived from the analysis, present a stark contrast to the extensive effort required for physical experiments.

The application of LazyNet to real-world data not only corroborates these findings but also facilitates the formulation of experimental plans. In this instance, the findings suggest that subsequent experiments should focus specifically on the Myd88 pathway rather than considering a broader range of possibilities, thereby optimizing laboratory resources. Through the integration of ScVelo and LazyNet, we have enhanced the predictive capabilities and temporal resolution of scRNA-seq analyses, establishing a pioneering approach in the field.

## Discussion

LazyNet integrates unique logarithmic and exponential paired layers to mimic mathematical operations, forming a robust ODE system. Across four case studies, LazyNet has demonstrated its proficiency in handling well-defined ODE systems and synthetic data. It has also offered some degree of interpretation of real-world data, achieving insights previously unattainable with conventional methods and handling volumes of data that were once considered impractical.

Despite these advancements, several limitations remain. Notably, the two real-world applications did not yield perfect solutions. This is often due to the physical experimental data having limited variability in treatments and temporal resolution, which restricts LazyNet’s applicability primarily to experimental modeling contexts. A potential solution is to utilize library-based experiments, such as deploying a library of inhibitors and recording temporal changes, or leveraging new pseudo-time compatible techniques like single-cell RNA sequencing and RNA velocity analysis. Other improvement directions include use better mechanism than log-exp pair mechanism and modularized network that can transfer learning.

Looking forward, LazyNet has the potential for broad applicability beyond biological systems. By incorporating three-dimensional coordinates, it could also be adapted for structural biology, and its principles could extend to fields like chemistry and materials science, where dynamic systems modeling is crucial.

## Methods

### Training

The training process commences by predicting the second data point from the first within each batch, and then iteratively progresses through the dataset in a batch-wise manner. Although a minimal batch size of two—focusing on changes between two consecutive data points—can suffice for basic predictions, larger batch sizes are generally adopted to enhance the overall accuracy of the model. The optimal batch size is carefully chosen to strike a delicate balance between capturing detailed data variations and mitigating the potential inaccuracies that smaller batches might introduce.

### Synthetic Data Studies

Both the HIV population and Gene Regulatory Network (GRN) studies utilized synthetic data. Details on the data and code used can be found in the Data and Code Availability section.

### Metabolite Data Study

Original literature predicts metabolomic outcomes based on proteomics data, the scope of the study was adjusted to predict phenotypic behaviors using the available metabolomics data. The dataset contains more than 100 sample data.

For this study, significant challenges related to data availability, with only 9 out of the 100 expected datasets being accessible. The proteomics data also not formatted and provided in matrix.

LazyNet was applied to analyze mass spectrometry data from eight engineered E. coli strains, focusing on their ability to synthesize three distinct biochemical products. There are 79 small molecules data in the dataset, and the network contains 80 inputs including one additional time variable. To refine our analysis, we expanded each dataset to 500 data points using the Savitzky-Golay filter.

Details on the data and code used can be found in the Data and Code Availability section.

### Single-Cell RNA Sequencing (scRNA-seq)

The scRNA-seq data is based on previous studies concerning monocyte dynamics activated by LPS. Details about the dataset and processing are provided in the Data and Code Availability section. This dataset posed significant challenges for LazyNet due to its real-world complexity, including inherent noise and a limited sample size of 6,000 cells across various time points, offering only a modest number of analyzable samples. Moreover, the necessity to manage an overwhelming number of genes—of which only a subset was relevant—significantly complicated the analysis process.

To mitigate these challenges, we utilized Seurat to focus on 717 genes identified through Principal Component Analysis (PCA), which aided in generating a robust Uniform Manifold Approximation and Projection (UMAP). The UMAP not only validated our predictions but also facilitated meaningful comparisons with prior studies. The cells were organized into seven clusters with pseudo-time points assigned, and the dataset was expanded to include 140 data points using the Savitzky-Golay filter for smoothing as the trainset for LazyNet. The goal was to extend the trajectory and uncover any clues generated by the prediction.

